# Hox11-expressing interstitial cells contribute to adult skeletal muscle at homeostasis

**DOI:** 10.1101/2022.06.14.496124

**Authors:** Corey G.K. Flynn, Qingyuan Guo, Paul R. Van Ginkel, Steven M. Hrycaj, Aubrey E. McDermott, Angelo Madruga, Deneen M. Wellik

## Abstract

Adult skeletal muscle possesses remarkable regenerative capacity. This is attributed to tissue-specific stem cells, satellite cells. Interstitial stromal cells also play critical roles in muscle, and we have previously reported that *Hoxa11* and *Hoxd11*, expressed in the interstitial cells of muscles that attach to the zeugopod (radius and ulna), are critical for the proper patterning and development of these muscles during embryogenesis. Using a Hoxa11eGFP knock-in reporter, we show that expression continues in a subset of muscle interstitial cells through adult stages. Using *Hoxa11-CreERT2* mediated lineage reporting induced at adult stages, we observe lineage initiation only in the interstitial cells of muscle, as expected. However, this Hoxa11-expressing interstitial cell lineage progressively contributes to muscle fibers at postnatal and adult stages. The contribution to these muscles at adult homeostasis significantly exceeds parallel *Pax7-CreERT2* mediated lineage labeling performed in parallel. To confirm that interstitial cell nuclear contents are contributed to muscle fibers, we additionally used the nuclear specific lineage reporter, *ROSA-LSL-H2BmCherry* with *Hoxa11-CreERT2* and observe that Hoxa11-expressing interstitial cells contribute their nuclei to myofibers. *Hox* lineage contribution is observed into all four muscle sub-types over months of lineage labeling. At no point after Hoxa11-mediated lineage induction do we observe lineage labeling into Pax7-expressing satellite cells. This adds to a small but growing body of evidence that supports a satellite cell-independent source of muscle tissue *in vivo*.

**Summary Statement:** Hoxa11 expression marks a novel population of muscle interstitial cells capable of extensive, satellite cell-independent contribution to skeletal muscle fibers during adult homeostasis.

## Introduction

Roles for *Hox* genes in the embryonic development of the skeletal system are well established, with loss-of-function of *Hox* paralog groups leading to dramatic homeotic transformations of the axial vertebrae, perturbed morphogenesis of the limb skeleton, and disrupted organogenesis. In limb skeletal patterning, *Hox* paralog groups *Hox9 – Hox13* have region specific function and loss of *Hoxa11* and *Hoxd11* gene function, dramatic mis-patterning of the radius and ulna is observed (Davis and Capecchi, 1996; Fromental-Ramain et al., 1996a; Fromental-Ramain et al., 1996b; Wellik and Capecchi, 2003). In compound *Hox11* mutants where three of the four *Hoxa11* and *Hoxd11* alleles are absent, no skeletal phenotype results, but patterning of the zeugopod-attached muscles and tendons is disrupted, supporting a direct role for *Hox11* genes in muscle patterning (Swinehart et al., 2013).

*Hoxa11* gene expression initiates broadly in the lateral plate mesoderm of early developing limb buds, but expression quickly localizes to the zeugopod region as it forms, and can be observed throughout development in the stromal cells surrounding the developing cartilage and bone, in the tendons, and in a continuous population of connective tissue/interstitial stroma surrounding the zeugopod-attached myofibers (Swinehart et al., 2013). In this report, we show that Hoxa11eGFP expression in zeugopod-attached muscle interstitial cells continues throughout postnatal and adult life.

Limb skeletal muscle progenitors derive from the embryonic somites where muscle precursors are marked by expression of Pax3 and, later in development, Pax7 (Chal and Pourquié, 2017; Sefton and Kardon, 2019). Pax3+ progenitors delaminate from the somites and migrate to the limb bud. These precursors undergo additional steps of myogenesis to generate multinucleated myocytes (Chal and Pourquié, 2017; Nassari et al., 2017; Sefton and Kardon, 2019). After maturation, muscles retain significant plasticity with the ability to hypertrophy with increased use and to repair in response to injury. Repair of skeletal muscle is known to rely on satellite cells, stem cells set aside inside of the basal lamina of multi-nucleate muscle fibers that are self-renewing and characterized by the expression of Pax7 (Chal and Pourquié, 2017; Lepper and Fan, 2010; Lepper et al., 2011; Pawlikowski et al., 2015; Yin et al., 2013). Ablation of Pax7-expressing satellite cells leads to a profound deficit in the ability of skeletal muscle to regenerate in response to injury (Lepper et al., 2011; Sambasivan et al., 2011). Pax7-mediated lineage studies have demonstrated that satellite cells contribute to adult skeletal muscles at homeostasis, however, ablation of satellite cells at adult stages, in the absence of injury, does not lead to early sarcopenia and, surprisingly, skeletal muscle remains capable of hypertrophy (Fry et al., 2015; Jackson et al., 2012; Keefe et al., 2015; McCarthy et al., 2011).

A heterogenous population of stromal cells, called interstitial cells, lie outside and surround the basal lamina of the muscle fibers. Interstitial cells are critical for embryonic muscle patterning; neither somite-derived muscle progenitors nor satellite cells possess intrinsic muscle patterning information (Aoyama and Asamoto, 1988; Duprez, 2002; Lance-Jones, 1988; Mathew et al., 2011; Michaud et al., 1997; Nassari et al., 2017). Multiple sub-populations of interstitial cells have been defined, including pericytes, PW1+/Pax7-interstitial progenitor cells (PICs), fibroblasts, fibroadipogenic progenitors (FAPs), and smooth muscle-mesenchymal cells (SMMCs) (Giordani et al., 2019; Malecova and Puri, 2012; Mathew et al., 2011; Mierzejewski et al., 2020a; Tedesco et al., 2017). How these interstitial cell populations are distinct from one another is not fully understood, though there is at least some overlap between most of these subsets (Malecova and Puri, 2012; Tedesco et al., 2017). Previous work has shown that interstitial stromal cells support myofiber function and regeneration at adult stages, and can impact satellite cell quiescence, activation, and migration (Heredia et al., 2013; Joe et al., 2010; Kuswanto et al., 2016; Murphy et al., 2011; Nassari et al., 2017; Tatsumi et al., 2009; Thomas et al., 2015; Tidball and Villalta, 2010; Uezumi et al., 2010).

A small but growing number of publications have reported that subsets of interstitial cells possess myogenic potential, but the *in vivo* myogenic potential for these cells remains controversial (Dellavalle et al., 2011; Doyle et al., 2011; Esteves de Lima et al., 2021; Giordani et al., 2019; Liu et al., 2017; Mierzejewski et al., 2020b; Mitchell et al., 2010; Qu-Petersen et al., 2002). Twist2-directed genetic lineage labeling from an interstitial cell sub-population demonstrated direct contribution to adult skeletal muscle fibers *in vivo* (Liu et al., 2017). Twist2-expressing cells were reported to contribute specifically to Type IIb/x myofibers at homeostasis and following injury. Recently, it was reported that early lateral plate lineage cells from a *Scx-Cre* reporter show contribution to muscle fibers and some of the satellite cell population specifically at the myotendinous junction during embryogenesis (Esteves de Lima et al., 2021).

In this study, we use our *Hoxa11eGFP* reporter and *Hoxa11-CreERT2* in combination with *ROSA26-LSL-tdTomato* and *ROSA26-LSL-H2BmCherry* lineage reporters to examine Hoxa11 expression and the lineage contribution from Hox-expressing cells from embryonic to adult stages (Blum et al., 2014; Madisen et al., 2010; Nelson et al., 2008; Pineault et al., 2019). We demonstrate that Hoxa11eGFP expression is restricted to a sub-population of zeugopod-attached, muscle interstitial cells at all stages of life. Flow cytometry and immunofluorescent analyses show that Hoxa11-expressing cells are a non-hematopoietic, non-endothelial, non-satellite cell stromal population with strong overlap with Twist2 and partial overlap with Tcf4 and PDGFRα. *Hoxa11-CreERT2* lineage induction from any stage results in initiation of labeling only in interstitial cells, as expected. Induction of lineage from E12.5 shows restriction of reporter expression to only the muscle interstitial cells throughout the remainder of embryonic development. However, at postnatal and adult stages, the Hoxa11 lineage begins to show contribution to myofibers. This contribution is progressive; reporter expression increases in skeletal muscle fibers in the weeks after induction. In parallel experiments comparing *Hoxa11-CreERT2* and *Pax7-CreERT2* to induce lineage labeling from the same *ROSA26-LSL-tdTomato* reporter, Hoxa11 lineage contribution is significantly higher than Pax7-expressing satellite cell contribution to forelimb muscles during adult homeostasis. Intriguingly, the extent of *Hoxa11*-mediated lineage labeling varies in a reproducible pattern in different muscle groups even though *Hoxa11*-expressing interstitial cells are present in all zeugopod-attached muscles. Contribution of *Hoxa11*-expressing interstitial cells to muscle fibers was further validated by using a *Hoxa11-CreERT2;ROSA-LSL-H2BmCherry* reporter. Using this reporter, we demonstrate that Hoxa11-expressing interstitial cells contribute their nuclei to myofibers. The Hoxa11 lineage does not mark Pax7-expressing satellite cells at any time point, supporting *Hox11*-expressing cell contribution to myofibers that is independent from the satellite cell pool. Taken together, these data support Hoxa11-expressing interstitial cells are a population of non-satellite cell muscle progenitors *in vivo*. This report adds to the growing evidence for novel, non-satellite cell sources of muscle progenitors *in vivo* in mammals.

## Results

### Hoxa11eGFP expression is maintained in skeletal muscle interstitial cells throughout life

During development, Hoxa11eGFP expression in muscle is restricted to the zeugopod-attached muscles, specifically in the non-endothelial interstitial stroma (Swinehart et al., 2013). We examined expression at embryonic, postnatal, and adult stages and observed that expression is maintained throughout life in zeugopod-attached skeletal muscle interstitial cells (**Fig. 1A**). Using our previously validated *Hoxa11-CreERT2* to induce recombination of a *ROSA26-Lox-STOP-Lox-tdTomato* lineage reporter (Hoxa11iTom; Madisen et al., 2010; Pineault et al., 2019), we observe that 3 days after a single 5mg tamoxifen dose, Hoxa11iTom cells are distributed throughout the interstitial cells of all forelimb muscles (**Fig 1B**). Hoxa11iTom lineage is induced at approximately 90% efficiency at this tamoxifen dose and overlaps with Hoxa11eGFP live reporter expression, validating our lineage reporter (**Fig 1C, Fig. S1**). Additionally, Hoxa11 expression in muscle is only in the interstitial cells (**Fig. S2**).

**Figure 1.**
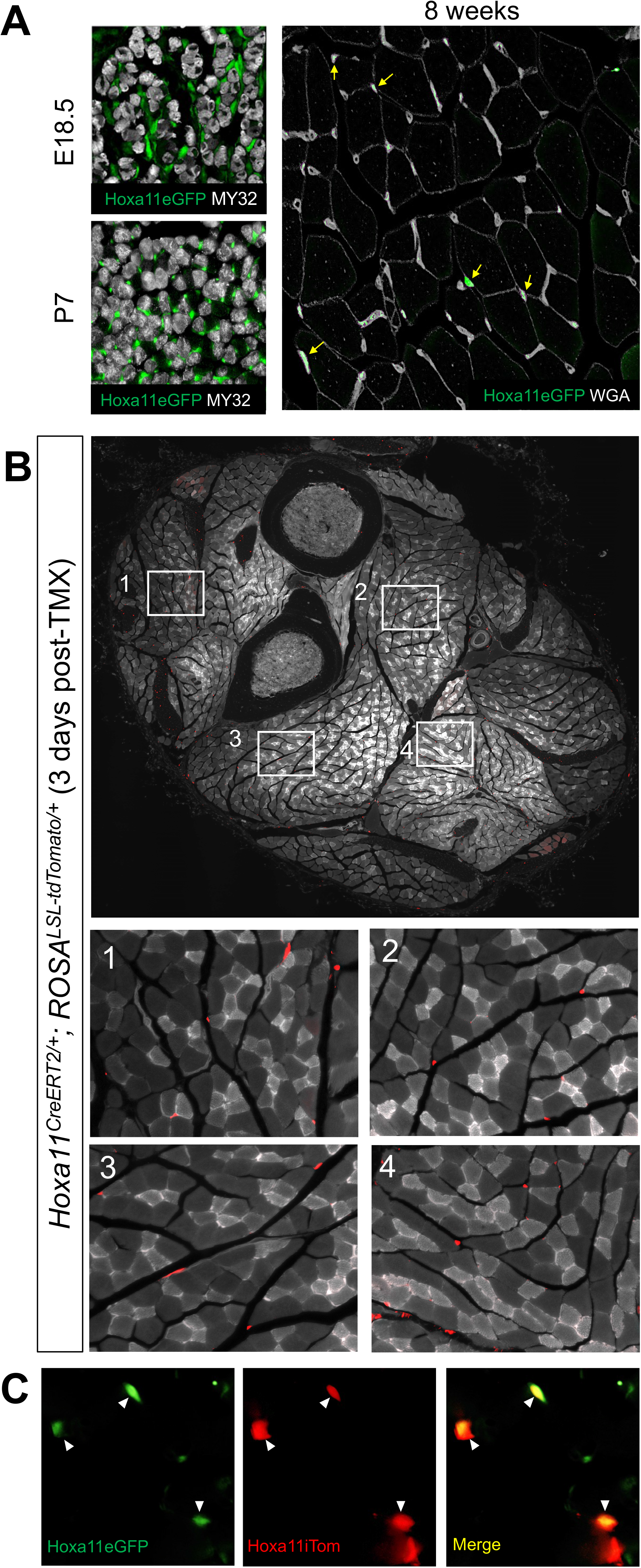
Hoxa11 expression is maintained in skeletal muscle interstitial cells throughout life. (**A**) Hoxa11eGFP real-time reporter shows expression (green) at E18.5 and P7 surrounding muscle fibers (MY32, white). At 8 weeks of age, Hoxa11eGFP (green, yellow arrows) is expressed in the interstitium visualized with WGA (white). (**B**) High magnification images from a whole forelimb cross section show tdTomato expression (Hoxa11iTom, red) in the interstitium throughout forelimb muscles. (**C**) Hoxa11 lineage reporter (Hoxa11iTom, red) overlaps with real-time reporter Hoxa11eGFP (green, white arrowheads) in interstitial cells 3 days post-tamoxifen.

### Hoxa11eGFP-expressing cells are a non-hematopoietic, non-endothelial, non-satellite cell sub-population of muscle interstitial cells

Hoxa11eGFP-expressing cells were assessed by flow cytometry to investigate how these cells correlate with previously described populations of mononuclear cells present in skeletal muscle. Live, mononucleated cells were enzymatically isolated from limb muscles attached to the zeugopod (Liu et al., 2015), and cells were gated for CD45 and Ter119, to identify hematopoietic cells. Hoxa11GFP-positive cells were essentially absent from the blood lineage, as expected (**Fig 2A**). Non-hematopoietic cells were gated for CD31, an endothelial cell marker, and we observed that Hoxa11GFP-positive cells were also largely absent from the endothelial cell population (**Fig. 2B**). Of the non-hematopoietic, non-endothelial population, approximately 2% of remaining cells were Hoxa11eGFP-positive. Cells were further gated to identify satellite cells, FAP cells and SMMCs (Smooth Muscle-Mesenchymal Cells) using the Sca1, VCAM1 and ITGA7 cell surface markers (Giordani et al., 2019; Joe et al., 2010; Liu et al., 2013). While most non-hematopoietic, non-endothelial cells are Sca1-negative (>75%), more than 50% of Hoxa11GFP-positive cells segregated as Sca1-positive (**Fig. 2C**). Within this Sca1^+^/Hoxa11eGFP^+^ population, the majority (91%) of Hoxa11eGFP-positive cells are Itga7-negative, correlating with previously identified FAPs (CD45+/CD31-/Sca1+/Itga7+, (Joe et al., 2010)). Among the non-hematopoietic, non-endothelial, Hoxa11eGFP-positive cells that are Sca1-negative (just under 50% of total Hoxa11eGFP population), additional labeling with ITGA7 and VCAM1 showed that the VCAM1+/Itga7+ satellite cell population contains essentially no Hoxa11eGFP+ cells (**Fig. 2E**). A small fraction of the total Hoxa11eGFP+ cell population (^~^7%) segregated as CD45-/CD31-/Sca1-/Itga7+/VCAM1-, markers reported recently to be a sub-population of interstitial SMMCs that possess myogenic potential (Giordani et al., 2019).

**Figure 2.**
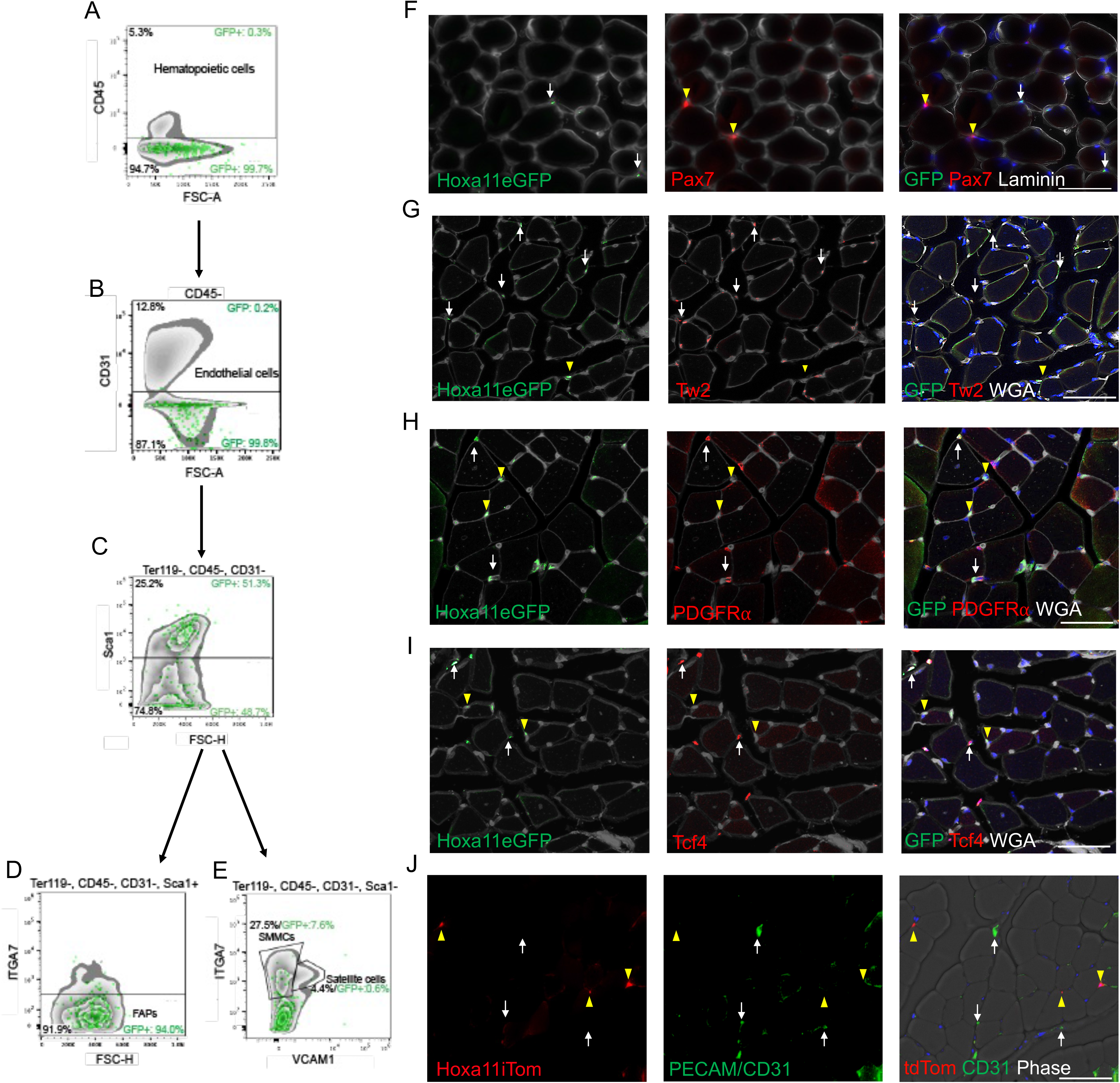
Hoxa11eGFP-positive cells are a subset of interstitial cells. Flow cytometric analyses of mononucleated live cells from zeugopod muscles of 14-week-old mice stained with CD45, TER119, CD31, Sca1, Itga7 and VCAM1. After sorting by forward scatter, side scatter and using DAPI for exclusion of dead cells, Hoxa11eGFP+ cells sort as non-hematopoietic (CD45- and TER119-negative), (**A**), and non-endothelial (CD31-negative), (**B**). (**C**) Within the non-hematopoietic, non-endothelial population, Hoxa11eGFP+ cells sort as 51% Sca1-positive and 49% Sca1-negative cells. (**D**) The majority of Hoxa11eGFP+/Sca1+ cells are Itga7-negative (FAPs). (**E**) The Hoxa11eGFP+/Sca1-population is negative for VCAM1, and Hoxa11eGFP+/Sca1-cells do not sort with the Itga7+/VCAM+ satellite cells. Approximately 8% of the Hoxa11eGFP+/Sca1-cells are Itga7+/VCAM-, a marker combination used to identify SMMCs. Hoxa11eGFP+ cells are represented as green dots on top of gray-scale density plots of non-GFP labeled cells. Forelimb ventral muscle sections of 8–10-week-old animals were immunostained for GFP, Pax7 and WGA (**F**); GFP, Twist2 and WGA (**G**); GFP, PDGFRα and WGA (**H**); GFP, Tcf4 and WGA (**I**). (**F**) Hox11eGFP+ (arrows) cells do not overlap with Pax7+ satellite cells (arrowheads). Arrows (white) in **G-J** indicate double positive cells. Arrowheads (yellow) in **G-J** indicate Hox11eGFP+ cells. (**J**) Hoxa11 lineage positive cells (Hoxa11iTom, arrows), 4-days post tamoxifen, do not colocalize with PECAM/CD31(arrowheads). Scale bars, 50 μm.

We further characterized Hoxa11eGFP-expressing cells *in situ* by co-immunofluorescent immunohistochemistry. Hoxa11eGFP expression is observed in interstitial cells outside the basal lamina of myofibers and shows no overlap with satellite cells based on Pax7 and laminin immunostaining (**Fig. 2F**). The transcription factor, Twist2, is expressed in interstitial cells that possess previously reported myogenic potential (Li et al., 2019; Liu et al., 2017); we observe that most Hoxa11eGFP-expressing cells also express Twist2, though the two populations are not completely overlapping (**Fig. 2G**). Within the interstitial population, Hoxa11eGFP also shows partial overlap with PDGFRα, a marker used to identify FAPs, and low overlap with Tcf4, a connective tissue fibroblast marker (Joe et al., 2010; Mathew et al., 2011; Murphy et al., 2011; Uezumi et al., 2010; Uezumi et al., 2011). Using our *Hoxa11* lineage reporter (Hoxa11iTom) we further demonstrate that, *Hoxa11* lineage labeled cells (3-days after tamoxifen treatment) do not colocalize with PECAM/CD31^+^ endothelial cells (**Fig.2J**). These results corroborate and extend flow cytometry analyses and demonstrate that Hoxa11eGFP is expressed in a sub-population of muscle interstitial cells.

### *Hoxa11* lineage in muscle tissue remains restricted to interstitial cells during embryonic stages but begins contributing to myofibers at postnatal stages

We next sought to examine the lineage of the Hoxa11-expressing population. To assess embryonic lineage, pregnant dams were dosed with tamoxifen at E12.5 and resulting *Hoxa11^CreERT2/+^; ROSA^LSL-tdTomato/+^* embryos were collected at E18.5. Hoxa11iTom expression is observed exclusively in the interstitial and connective tissue cells of developing muscles (**Fig 3A**).

**Figure 3.**
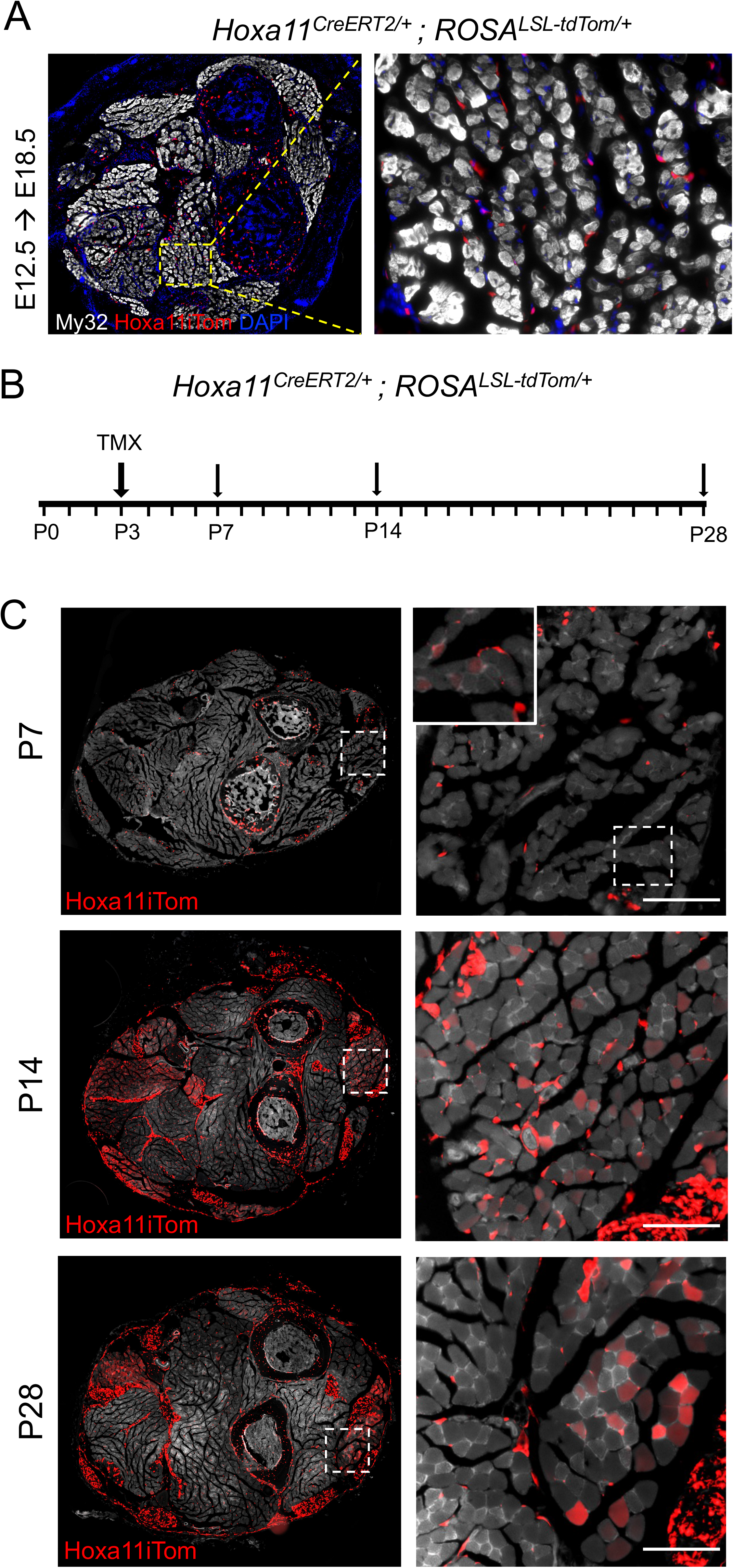
Embryonic *Hoxa11* lineage is exclusively interstitial; postnatal lineage begins to contribute to myofibers. At E12.5 the pregnant dams were dosed with 5mg Tamoxifen by intraperitoneal injection and embryos were collected at E18.5. (**A**) Representative cross-section of E18.5 forelimb stained for My32 (white) with Hoxa11iTom (red) visible in connective tissue and bone; nuclear staining shown with DAPI (blue). High magnification of E18.5 forelimb muscle shows tdTomato signal dose not overlap with My32. (**B**) At postnatal day 3 Hoxa11^CreERT2/+^; ROSA^LSL-tdTom/+^ mice were given 0.25mg Tamoxifen by intragastric injection and collected at P7, P14, and P28. (**C**) Full forelimb cross sections show tdTom (Hoxa11iTom) expression throughout forelimb tissues and high magnification images show clear tdTom expression in myofibers at P14 and P28. Inset of P7 muscle shows some faint tdtom+ myofibers when imaged at a higher exposure. Scale bars, 100μm

Postnatal lineage was examined by dosing *Hoxa11^CreERT2/+^; ROSA^LSL-tdTomato/+^* animals with tamoxifen on postnatal day 3 (P3) and collecting them at P7, P14, and P28 (**Fig 3B**). Whole forelimb cross-sections and higher magnification images show tdTomato expression within tendon, bone, and muscle at all time points. Hoxa11iTom is expressed throughout the zeugopod forelimb in muscle interstitial cells at P7. At this time point, very low levels of tdTomato expression can be observed in a small number myofibers. Images of P7, P14, and P28 taken at the same exposure show increasing number and intensity of tdTomato+ myofibers with time after induction.

A recent report identified an unexpected contribution to the myogenic lineage from embryonic lateral plate mesoderm using both quail-chick chimeras and using a *Scleraxis-Cre* allele and reported *Scx-Cre*-mediated contribution to the embryonic myogenic lineage at or near the myotendinous junction (Esteves de Lima et al., 2021). We observe no such localization to the myotendinous junction region at embryonic or adult stages (**Fig. S3**). Together our data supports a Hoxa11 lineage that is restricted to the muscle interstitium throughout embryonic stages but begins to progressively contribute to myofibers during the postnatal period.

### *Hoxa11*-expressing cells make progressive contribution to adult skeletal myofibers at homeostasis to a greater extent than *Pax7* lineage-induced satellite cells

To assess Hoxa11 lineage behavior at adult stages, *Hoxa11^CreERT2/+^; ROSA^LSL-tdTomato/+^* animals were dosed with 5 mg tamoxifen at 8 weeks of age and analyzed at 2-, 4-, 7-, 14-, and 56-days after tamoxifen induction (**Fig. 4A**). At 4 days post-induction, there is extensive lineage labeling of muscle interstitial cells and only faint tdTomato signal in a few muscle fibers. At continuing time points through 8 weeks post-induction, lineage contribution to muscle fibers progressively increases as seen in whole forelimb cross-sections and magnifications of three different muscles (**Fig. 4B**). Continual, progressive, and more extensive lineage labeling is observed at later stages (**Fig. S4**). The extent of lineage labeling into each muscle group varies, but with high reproducibility between animals. In Hoxa11iTom animals at 8 weeks after induction, the Extensor Carpi Ulnaris (ECU) shows 95.2% +/- 2.9 tdTomato-positive fibers. The Flexor Digitorum Profundus (FDP) shows 51.2% +/- 6.9 tdTomato-positive muscle fibers, and the Flexor Digitorum Sublimis (FDS) has 20.5% +/- 4.9 tdTomato-positive fibers. Despite differential lineage labeling into muscle groups, Hoxa11iTom-labeled interstitial cells are observed relatively uniformly throughout forelimb muscle tissue (**Fig. 1B; Fig S2**). Hindlimbs from these animals were also examined and the extent of lineage contribution to myofibers is comparable to the forelimb (**Fig. S5**).

**Figure 4.**
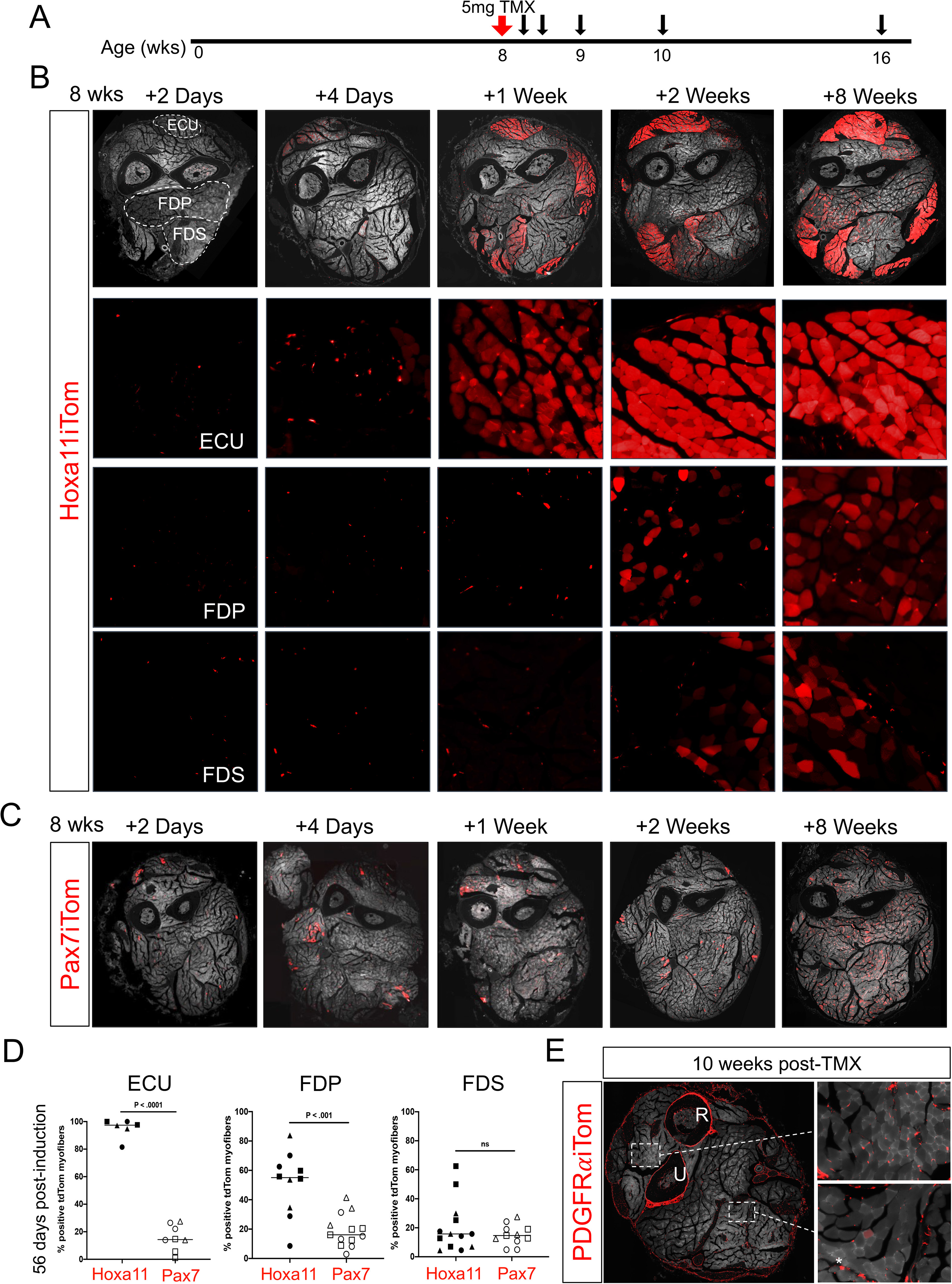
Hoxa11-expressing skeletal muscle interstitial cells progressively contribute to myofibers in the adult mouse forelimb. (**A**) *Hoxa11^CreERT2/eGFP^;ROSA^LSL-TdTom/+^* (Hoxa11iTom) mice were given a single intraperitoneal injection of 5 mg tamoxifen at 8 weeks of age and collected at 2 days, 4 days, 1 week, 2 weeks, and 8 weeks after tamoxifen induction. (**B**) Whole forelimb cross-sections were collected at time points indicated. Images of Extensor Carpi Ulnaris (ECU), Flexor Digitorum Profundus (FDP), and Flexor Digitorum Sublimis (FDS) muscles from Hoxa11iTom muscle show increasing number of tdTomato expressing myofibers over time. (**C**) Full forelimb cross-sections from *Pax7^CreERT2/+^; ROSA^LSL-TdTom/+^* (Pax7iTom) animals shows fewer tdTomato positive myofibers compared to Hoxa11iTom animals at the same time points. (**D**) Quantification of percentage tdTomato-positive myofibers from ECU, FDS, and FDP muscles from Hoxa11iTom and Pax7iTom animals 8 weeks post-induction (n=3 animals, 2-5 fields of view per muscle group analyzed). Statistics by Student’s t-test. ns, not significant. White dashed line marks borders of muscles shown in **B** and quantified in **D**, position of radius and ulna marked by R and U, respectively. (**E**) *PDGFRα^CreERT2/+^; ROSA^LSL-tdTomato/+^* (PDGFR*α*iTom) animals were treated with 5 mg tamoxifen at 8 weeks of age and collected 10 weeks later. Whole cross section of the forelimb shows tdTomato expression in connective tissues, bone tissue, tendons, and the muscle interstitium. High magnification images of skeletal muscle shows PDGFR*α*iTom expression in the muscle interstitium. A few myofibers were noticed to have low tdTomato expression marked by asterisks (*).

Parallel experiments were performed with the same reagents using *Pax7^CreERT2/+^; ROSA^LSL-tdTomato/+^* mice (Pax7iTom) (Murphy et al., 2011) to compare Hoxa11 lineage contribution to satellite cell contribution at homeostasis. Animals were given the same tamoxifen treatment and this resulted in ^~^90% efficient recombination in the *Pax7^CreERT2/+^; ROSA^LSL-tdTomato/+^* animals as well (**Fig. S1**). The observed lineage contribution of Pax7-expressing satellite cells was significantly less than from Hoxa11-expressing cells (**Fig. 4B-D**). Of note, Pax7 lineage contribution was more evenly distributed throughout the muscle groups, with low but consistent increases in the amount of lineage-labeled myofibers over time, consistent with findings reported previously in the hindlimb (Keefe et al., 2015; Pawlikowski et al., 2015). In examining the same three muscle groups, Pax7iTom contribution is 15.7% +/- 3.3 in the ECU, 21.5% +/- 3.1 in the FDP, and 16.4% +/- 2.1 in the FDS. High-magnification images of the ECU, FDP, and FDS muscle from Pax7iTom animals are shown in **Fig. S6**. Comparative quantification of lineage-positive myofibers of the FDP, ECU and FDS from both Hoxa11iTom and Pax7iTom animals at 8 weeks after induction shows Hoxa11iTom contribution is significantly higher in both the FDP and ECU muscles (**Fig. 4D**).

Our results show many similarities to those reported for Twist2-expressing interstitial cells (Liu et al., 2017) and, intriguingly, Hoxa11eGFP and Twist2 expression are highly overlapping. We sought next to interrogate whether this cellular lineage behavior occurs from other populations of muscle interstitial cells. PDGFRα is a more broadly expressed interstitial cell marker, so we utilized a *PDGFRα-CreERT2* and the *ROSA-LSL-tdTomato* reporter (PDGFRαiTom) to investigate the lineage contribution to myofibers (Rivers et al., 2008). Using this model, we do not observe lineage labeling into myofibers, even 10 weeks after adult induction (**Fig. 4E**). Of note, there is previously reported evidence that tdTomato mRNA can be packaged in extracellular vesicles that enter the myofiber, resulting in fusion-independent lineage reporter expression within muscle fibers (Murach et al., 2020). Close examination of the PDGFRαiTom shows a small number of myofibers (^~^5 in a whole forelimb cross-section) with low levels of red fluorescence, perhaps indicative of this type of non-specific labeling (**Fig. 4E**). However, the relative lack of myofiber lineage labeling in the PDGFRαiTom animals supports that lineage contribution in the Hoxa11iTom model does not result from leaky expression or fusion of cytoplasmic vesicles.

### *Hoxa11* lineage demonstrates nuclear contribution to myofibers

To provide additional evidence that the observed Hoxa11iTom lineage represents contribution of interstitial cells to myofibers and is not due to transport of tdTomato+ cytoplasm or tdTomato mRNA into muscle fibers, we generated a *Hoxa11*-dependent nuclear lineage reporter, by crossing our *Hoxa11-CreERT2* line with *ROSA-LSL-H2BmCherry* (Blum et al., 2014). Resulting *Hoxa11^CreERT2/+^; ROSA^LSL-H2B-mCherry/+^* (Hoxa11iH2BmCherry) animals were given 5mg tamoxifen at 6 weeks of age and collected at 4 days and 2 weeks after induction (**Fig 5A**). H2BmCherry-labeled myonuclei are clearly observed in muscle interstitial cells 4 days after tamoxifen induction (**Fig 5B**). By 2 weeks after lineage induction, Hoxa11iH2BmCherry+ nuclei can be additionally visualized within myofibers under the basal lamina (**Fig. 5C**). Hoxa11iH2BmCherry-positive nuclei were confirmed with DAPI staining (Fig. S7). These experiments provide strong support for *Hoxa11*-expressing interstitial cells contributing full cellular contents, including their nuclei, to muscle fibers *in vivo*.

**Figure 5.**
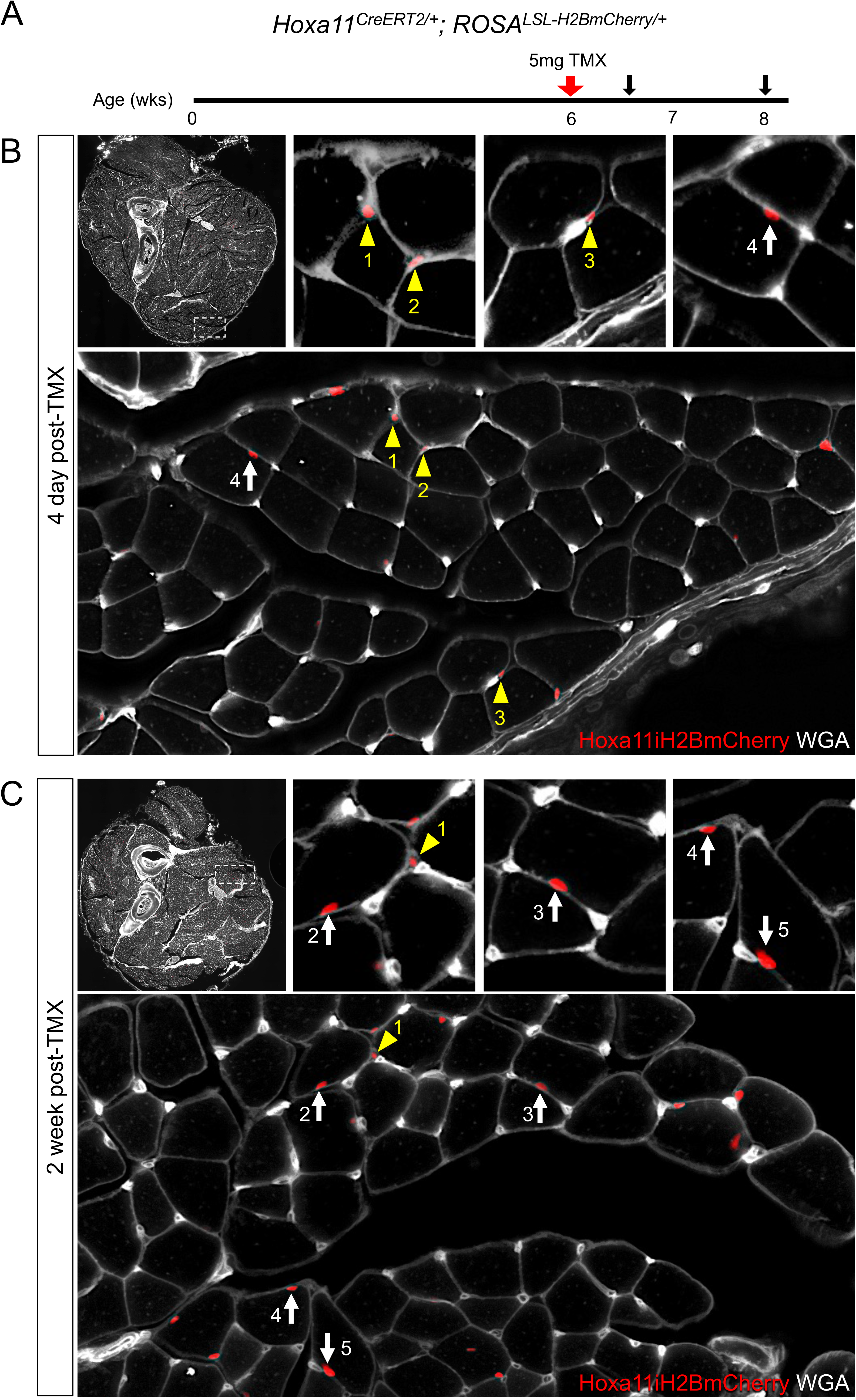
Hoxa11-expressing interstitial cells contribute their nuclei to myofibers. (**A**) *Hoxa11^CreERT2/+^; ROSA^LSL-H2BmCherry/+^* mice were given 5mg of tamoxifen at 6 weeks of age and collected 4 days and 2 weeks after dosing. (**B**) Top, left image shows a cross section of the forelimb of an adult mouse 4-days post-tamoxifen dosing. The larger, bottom image is a higher magnification image of the tissue in the white dashed box. The three top, right images are high resolution pictures of interstitial or myofiber nuclei from the bottom panel. (**C**) Top, left image shows a cross section of the forelimb of an adult mouse 2-weeks post-tamoxifen dosing. The larger, bottom image is a higher magnification image of the tissue in the white dashed box. The three top, right images are high resolution pictures of interstitial or myofiber nuclei from the bottom panel. Yellow arrowheads point to H2BmCherry labeled interstitial cell nuclei; white arrows point to H2BmCherry labeled myonuclei in **B** and **C**.

### Hoxa11 lineage contributes to all muscle fiber types

Previous publications have reported that *Twist2*-mediated contribution to muscle exhibits fiber-type specificity, contributing to only Type IIb/x myofibers (Li et al., 2019; Liu et al., 2017). Of note, these experiments were conducted on animals that were lineage labeled for up to four months. Observation of the Hoxa11iTom lineage in the hindlimb at 3 weeks following adult induction supports this specificity as almost no lineage labeling is observed in the soleus, which is comprised of mainly Type 1 myofibers while fairly extensive labeling is observed in the adjacent gastrocnemius, which is mainly Type IIb and IIx (**Fig. S5**)(Burkholder et al., 1994). However, upon evaluation of myofiber type specificity in Hoxa11iTom lineage at 8 months after induction of the reporter (10 months of age), Hoxa11 lineage contributes to all four muscle types, Type I, IIa, IIb, and IIx, is observed (**Fig 6A-D**).

**Figure 6.**
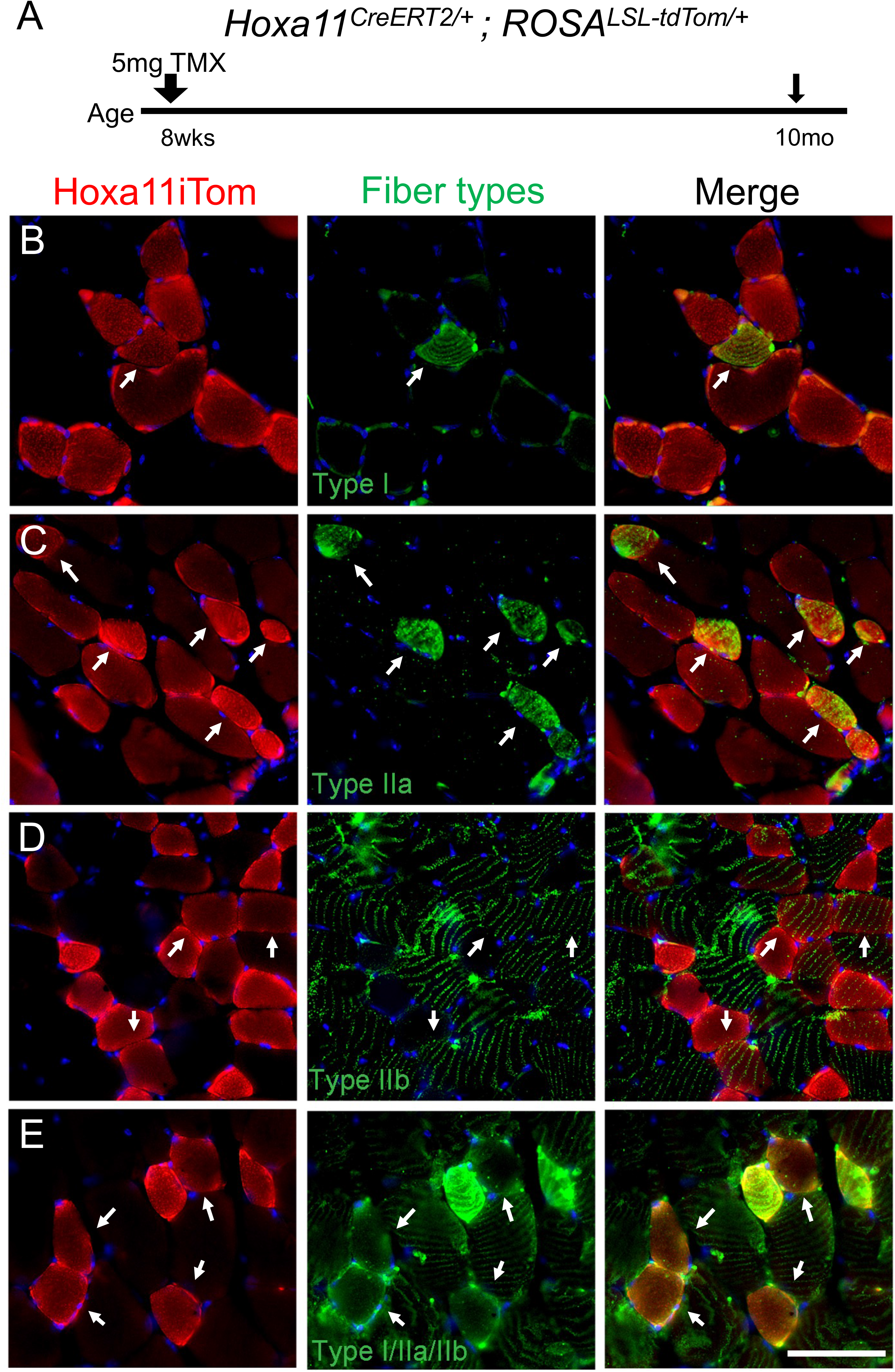
Hoxa11 lineage contributes to Type I, Type IIa, Type IIb, and Type IIx myofibers. (**A**) Mice were treated with tamoxifen at 8-weeks of age and collected at 10 months of age. (**B-D**) Hoxa11 lineage labeled myofibers are identified by tdTomato signal (Hoxa11iTom). Myofiber sub-types were identified by immunofluorescent staining with anti-myosin (Slow), anti-SC-71, and anti-BF-F3 (additional antibody details are available in Supplemental Table 1). (**E**) Type IIx fibers were identified by the absence of combined markers. Fibers shown are from the Gastrocnemius/Plantaris muscles. Lineage labeling is observed in all myofiber sub-types (white arrows). Scale bar = 100μm.

### Satellite cells are not lineage labeled by the *Hoxa11* lineage reporters

To determine whether Hoxa11-mediated lineage labeling results in reporter expression in satellite cells, we carefully examined co-expression of Pax7 and Hoxa11iTom in both the forelimb and the hindlimb of Hox11iTom animals. In the forelimb at 2 weeks after lineage induction, the ECU, FDP, and FDS were assessed for overlap of Hox11iTom and Pax7; we observed 0 instances of co-expression of Pax7 and Hoxa11iTom among a total of 1,048 individual, DAPI-stained cells analyzed (**Fig. 7A-D**). We further assessed Hoxa11 lineage and Pax7 antibody staining 8 weeks after lineage induction and again found no tdTomato+ satellite cells from 311 cells counted (**Fig. S8**). In the hindlimb, we analyzed three muscle groups, the Gastrocnemius/Plantaris, Soleus, and Tibialis Anterior for possible Hoxa11iTom and Pax7 co-expression. Consistent with data collected for the forelimb, no overlap of Hoxa11 lineage and Pax7 was observed out of 695 cells analyzed (**Fig. 7E-G**). Using our nuclear specific reporter (Hoxa11-H2BmCherry), we assessed whether any Pax7+ cells were H2BmCherry+. We observed zero Pax7+/mCherry+ cells 2 weeks after reporter induction (**Fig. 7H**) These results indicate Hoxa11-expressing cells do not contribute to or non-specifically label the satellite cell population.

**Figure 7.**
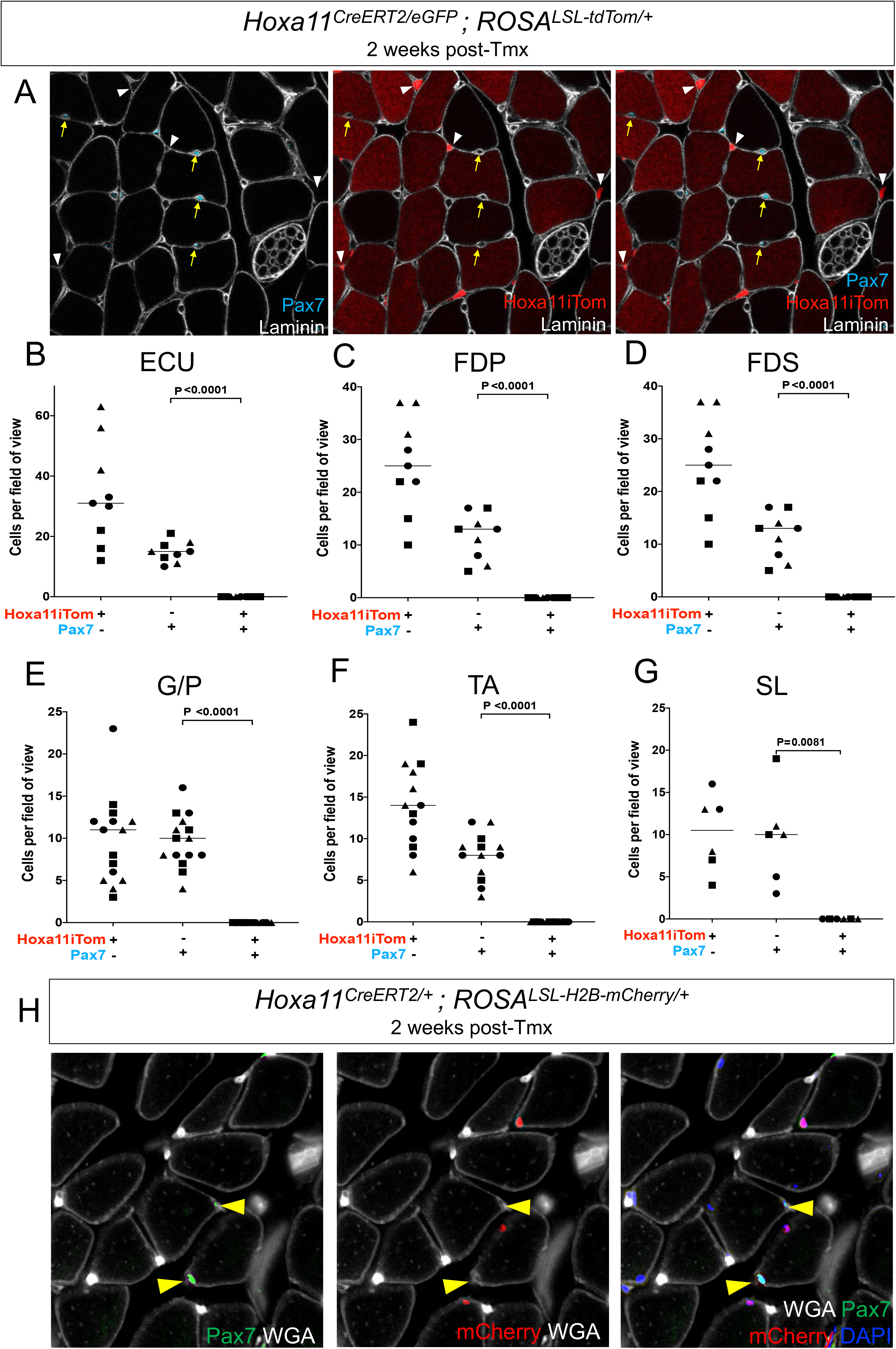
*Pax7*-expressing satellite cells are not lineage labeled by Hoxa11 lineage. *Hoxa11^CreERT2/+^; ROSA^LSL-tdTom/+^* mice given a single 5mg dose of tamoxifen by intraperitoneal injection at 8 or 12 weeks of age were collected 2-weeks later. (**A**) Pax7 cells (cyan, yellow arrows) are visualized under the basal lamina (white) with zero incidences of Hoxa11 lineage labeling (red) of Pax7-expressing cells in forelimb skeletal muscle at 2 weeks post-induction. (**B-G**). The number of Hoxa11iTom and Pax7 single- or double-positive cells were quantified in the Extensor Carpi Ulnaris (ECU), Flexor Digitorum Profundus (FDP), and Flexor Digitorum Sublimis (FDS) muscles of the forelimb and the Gastrocnemius/Plantaris (G/P), Soleus (SL), and Tibialis Anterior (TA) muscles of the hindlimb. (**H**) *Hoxa11^CreERT2/+^; ROSA^LSL-H2B-mCherry/+^* animals were given a single 5mg dose of tamoxifen by intraperitoneal injection at 6 weeks of age and immunostained for Pax7+ satellite cells 2 weeks after dosing; no overlap of Hoxa11iH2BmCherry and Pax7 were observed.

## Discussion

Uncovering the cell types that are responsible for muscle maintenance and growth is crucial to understanding skeletal muscle’s dynamic nature and its remarkable regenerative capacity. In this study we show definitive interstitial cell contribution to myofibers *in vivo* beginning at postnatal stages through genetic lineage labeling of Hoxa11-expressing interstitial fibroblasts. The Hoxa11 lineage initiates only in interstitial cells and remains solely in this population through embryonic development, but then contributes progressively to skeletal muscle fibers at postnatal and adult stages. Hoxa11 is not expressed in the satellite cell population. Further, we show that Hoxa11-expressing interstitial cells do not lineage label into satellite cells after lineage induction, indicating interstitial cell contribution is not mediated through the satellite cell pool. Thus, Hoxa11-expressing cells are a unique set of progenitors.

There has been mounting evidence that some skeletal muscle interstitial cells have myogenic potential. Studies at postnatal time points have indicated that PICs can contribute to muscle fibers upon transplantation and can differentiate into myocytes *in vitro*, however, their endogenous biological role remains unclear (Mitchell et al., 2010). Using a *Alkaline Phosphatase-CreERT2*, authors report that pericytes contribute to the satellite cell pool and to muscle fibers, but after the postnatal stages, researchers observed very few lineage labeled fibers, even following cardiotoxin injury (Dellavalle et al., 2011). Twist2-directed genetic lineage labeling of interstitial cells previously reported non-satellite cell contribution to myofibers at adult stages *in vivo* with many similarities to the findings in this report. The strong overlap between Twist2 and Hoxa11eGFP expression suggests these two transcription factors mark a subset of interstitial cells that are relevant for this activity *in vivo*. The absence of lineage labeling into muscle fibers using *PDGFRα-CreERT2*, which is a broader, less specific marker of interstitial cells, further supports the notion that only of a unique subset of interstitial cells is capable of contributing to myofibers *in vivo*.

Another fibroblast population with potential myogenic activity was recently reported using single cell RNA-sequencing in conjunction with mass cytometry and his new population was named smooth muscle mesenchymal cells (SMMCs). These cells are ITGA7+VCAM1-, thus distinct from satellite cells which are VCAM1+ (Giordani et al., 2019). SMMCs were reported to have myogenic capacity *in vitro* and to contribute to myofibers upon transplantation into injured muscle. Our flow cytometry data shows relatively few (^~^7%) Hoxa11eGFP+ cells in this ITGA7+VCAM1-population, however, more rigorous genetic and transcriptomic comparisons will serve to clarify the relationship between these cells.

The wealth of literature and decades of research on satellite cells and their critical role as myofiber stem cells puts an exceedingly high burden of proof on new biology that suggests there may be an additional, interstitial cell progenitor pool important during mature muscle homeostasis. Strong genetic evidence of cellular behavior *in vivo* was provided by the reports that Twist2-expressing interstitial cells contribute to muscle fibers at homeostasis (Li et al., 2019; Liu et al., 2017). In this report, we show very similar ROSA-tdTomato lineage contribution from an independently generated, distinct genetic locus of another transcription factor (Hoxa11) that is expressed in a similar subset of muscle interstitial cells. There is reported evidence of non-specific cytoplasmic transport of fluorescent reporter mRNAs and proteins to other cells (Murach et al., 2020). To address this potential caveat, we carried out an additional line of genetic experiments using the ROSA-LSL-H2BmCherry lineage reporter. Using this nuclear localized lineage reporter, we show initial labeling in interstitial cell nuclei, but relatively rapid and distinct contribution of H2BmCherry-labeled nuclei into muscle fibers. Further, we show that *PDGFRα-CreERT2-induced* ROSA-tdTomato lineage labeling, which results in extensive labeling of a broader population of interstitial cells and connective tissue, does not lead to contribution to muscle fibers even 10 weeks after induction (with the exception of a few faint red muscle fibers that may indeed be reflective of cytoplasmic content sharing). Thus, the collective evidence strongly supports the specificity and uniqueness of a subset of interstitial cells with myogenic potential *in vivo*.

Further investigation is needed to understand more about this unique and important subset of interstitial cell myofiber progenitors, including the degree of stemness they possess, their contribution to myofibers during aging, exercise, and in response to injury, and their behavior *in vitro* and in transplantation experiments. This understudied population and its ability to contribute to muscle fibers at homeostasis *in vivo* has exciting potential to lead to new therapies for those suffering from myopathies and muscular dystrophies.

## Methods and Materials

### Mouse models

Generation of mouse models *Hoxa11eGFP* (Nelson et al., 2008), *Hoxa11-CreERt2* (Pineault et al., 2019), *Pax7-CreERT2* (Murphy et al., 2011), *PDGFRα-CreERT2* (Rivers et al., 2008), and *ROSA-LSL-H2B-mCherry* have been previously described. The *ROSA26-CAG-loxP-STOP-loxP-tdTomato* (JAX stock no. 007909, Madisen et al., 2010) was purchased from The Jackson Laboratory. Creation of *Hoxa11iTom* mice was achieved by crossing *Hoxa11^CreERT2/+^* males with *Hoxa11^eGFP/+^; ROSA^LSL-tdTomato/LSL-tdTomato^* females. Generation *Pax7iTom* mice was achieved by crossing *Pax7^CreERT2/+^* males with *ROSA^LSL-tdTomato/LSL-tdTomato^* females. All mouse colonies were maintained on a mixed genetic background. Both male and female mice were used in all experiments (and all control and mutant pairs were sex-matched). Animals used in this study were euthanized by CO_2_ followed by cervical dislocation. All procedures described are in compliance with protocols approved by the University of Wisconsin-Madison and the University of Michigan Committee on Animal Care and Use Committees.

### Tamoxifen treatment

Adult mice were given a single intraperitoneal (IP) injection of 5mg tamoxifen (Sigma T5648) dissolved in corn oil at 8 weeks of age and collected at indicated time points. Embryonic induction was achieved by mating *Hoxa11^CreERT2^*^/+^ males to *Hoxa11^eGFP/+^; ROSA^LSL-tdTomato/LSL-tdTomato^* females and presence of the vaginal plug was checked each morning. Pregnant dams were given 2mg tamoxifen and 1mg/mL progesterone dissolved in corn oil via intraperitoneal injection at indicated embryonic time points (Pineault et al., 2019). Postnatal animals were given 0.25mg of tamoxifen dissolved in corn oil via intragastric injection on postnatal day 3.

### Tissue Preparation

Left and right zeugopods were collected. Skin and soft tissues were removed from the region and the zeugopod muscles and skeleton was isolated. Muscles were either carefully dissected off the bone or muscle was left attached to the bone. Dissection was done in PBS. Muscle groups and intact limbs were fixed in 4% paraformaldehyde (PFA, Sigma) in PBS shaking at 4 °C for 1-3 days. Intact limbs were decalcified in 14% ethylenediaminetetraacetic acid (EDTA, EMD Millipore) for 3-5 days (postnatal animals) or 6-7 days (adults) at 4 °C. Samples were cryoprotected in 30% sucrose in PBS overnight before embedding in OCT compound (Fisher, cat no. 4585). Tissue was stored a −80 °C. Cryosections of 10-14 μm were analyzed by IF.

### Immunohistochemistry

Sections were rehydrated with PBS. In some cases, antigen retrieval was performed by heating slides in a citrate buffer for 10-30 min and blocked with TNB buffer (Perkin Emler FP1020). Sections were incubated in primary antibodies overnight at 4 °C and then incubated in secondary antibodies for 2 hours at room temperature and counterstained with DAPI for nuclear visualization. Antibody information is provided in supplementary information (**Table S1**). tdTomato and mCherry were visualized directly with no antibody staining. Slides were mounted with ProLong Gold antifade reagent (Invitrogen, cat no. P36930). Staining using mouse monoclonal antibodies utilized a mouse-on-mouse immunodetection kit (Vector Labs, cat no. BMK-2202) according to vendor instructions.

Imaging was performed on the Nikon eclipse Ti, Keyence BZ-X800, and Leica SP8 3X STED Confocal. Image editing was preformed using ImageJ and Photoshop, and larger images were stitched together (when necessary) on photoshop or by the Keyence BZ-X800 analyzer.

### Quantification

For calculation of percent positive tdTomato myofibers from Hoxa11iTom and Pax7iTom mice, multiple representative fields of view, from the ECU, FDP, and FDS muscles collected 8 weeks after Cre induction, were complied. A blinded participant counted the total number of myofibers in the field of view and the total number of tdTomato positive. Percent of tdTomato positive myofibers was calculated; data was transferred to GraphPad Prism9 where statistical analysis for significance, between Hoxa11iTom and Pax7iTom ECU, FDP, and FDS muscles, and standard error was performed. To assess overlay of Hoxa11 lineage labeled cells with Pax7, tissue from Hoxa11iTom mice was collected at 2 weeks and 8 weeks post induction; sections were stained using an anti-Pax7 antibody (DSHB). 3-5 fields of view from each animal (n=3 for both forelimb and hindlimb analyses) were used; Pax7 and Hoxa11iTom cells were identified, counted, and profiled into three groups: Hoxa11iTom+/Pax7-, Hoxa11iTom-/Pax7+, or Hoxa11iTom+/Pax7+. Statistical analyses were carried out by an unpaired Student’s t-test (GraphPad Prism9).

### Flow Cytometry

To isolate mononuclear populations from muscle, zeugopod-attaching muscles were removed from the forelimb and hindlimb. Dissected muscles were chopped into smaller pieces followed by digestion in 700 U/mL collagenase type II (Gibco, cat no. 17101-015) at 37°C for 1 hour. After spin and trituration steps muscles underwent a second digestion in 100 U/mL collagenase type II and 1.1 U/mL dispase (Gibco, cat no. 17105-041) for 30 min at 37°C. Cells were washed, triturated, spun, and filtered through a 40-micron cell strainer before being placed in 1xPBS with 2% FBS (Gibco). Flow cytometry experiments were carried out on the ThermoFisher Attune NxT Flow Cytometer BRYV. Markers and antibody information is provided in supplemental information (**Table S1**). Analysis was preformed using FloJo.

## Acknowledgments

We sincerely thank and are grateful for experiments performed by Anna P. Miller, PhD and assistance provided by Alex Hurley. We’d like to thank Gabrielle Kardon, PhD for providing the Pax7-CreERT2 mouse model and Barak Blum, PhD for providing the ROSA-H2B-mCherry mouse model, this work would not have been possible without those animals. Further, this work was supported by AR072511 (DMW) and the Flow Cytometry Laboratory at University of Wisconsin-Madison and their funding source CA014520.

---

**Figure S1.**
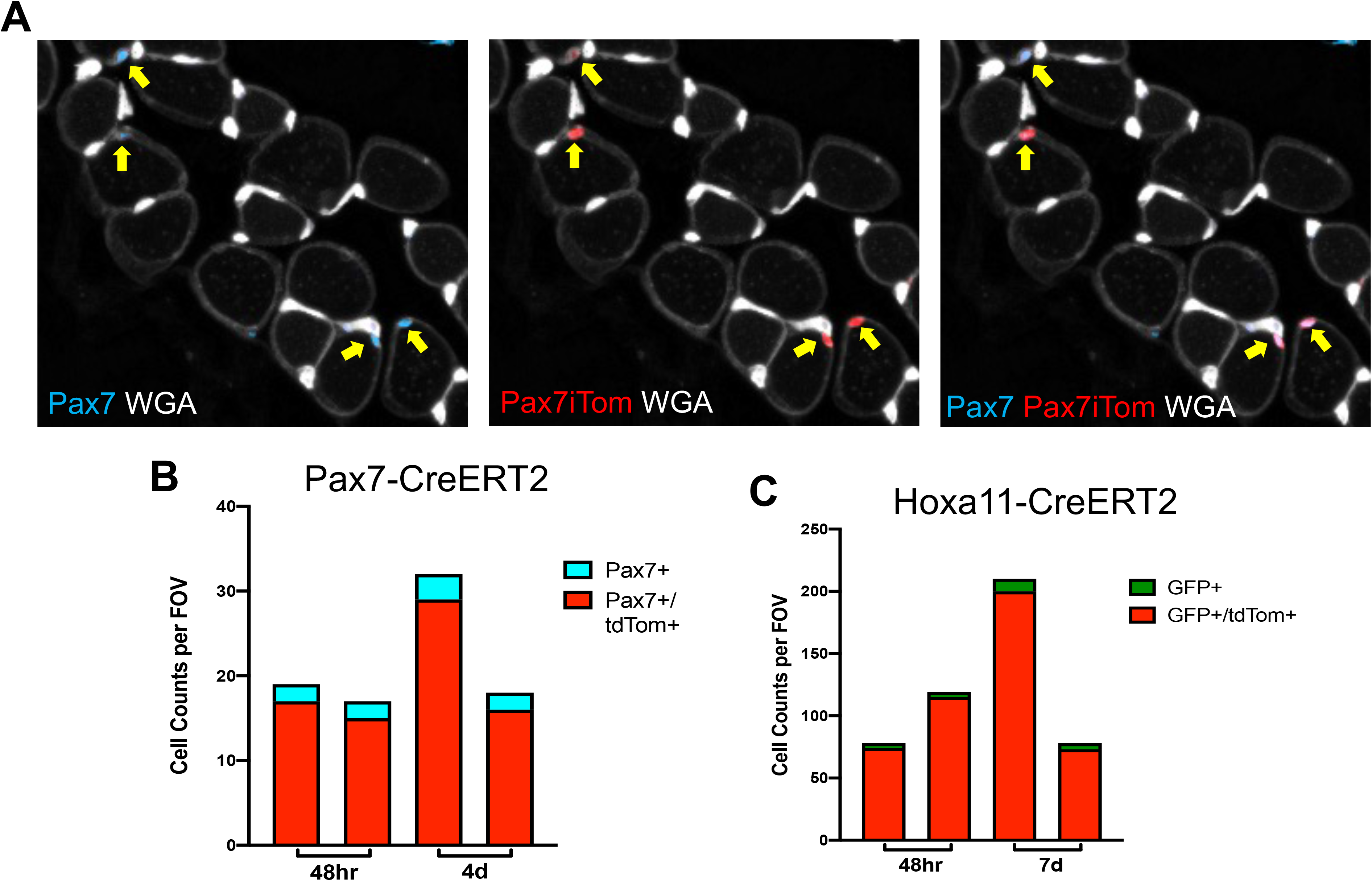
Both *Hoxa11-CreERT2* and *Pax7-CreERT2* show approximately 90% efficiency with lineage reporter ROSA-LSL-tdTomato. (**A**) Images show Pax7 IF staining (blue) overlaps with tdTomato (red) in Pax7 lineage reporter animals (Pax7iTom) 4 days after tamoxifen treatment. (**B**) Quantification of Pax7-CreERT2 efficiency was assessed by counting the number of Pax7 antibody-stained cells and tdTomato labeled cells at 48hrs (n= 2 animals) and 4 days (n= 2 animals) after Tamoxifen treatment. (**C**) Quantification of Hoxa11-CreERT2-induced recombination of ROSA-LSL-tdTomato with a single 5mg bolus of tamoxifen was assessed by counting the number of Hoxa11eGFP and tdTomato labeled cells at 48hrs (n=2 animals) and 7 days (n=2 animals) after tamoxifen administration. Both Cres resulted in approximately 90% efficiency.

**Figure S2.**
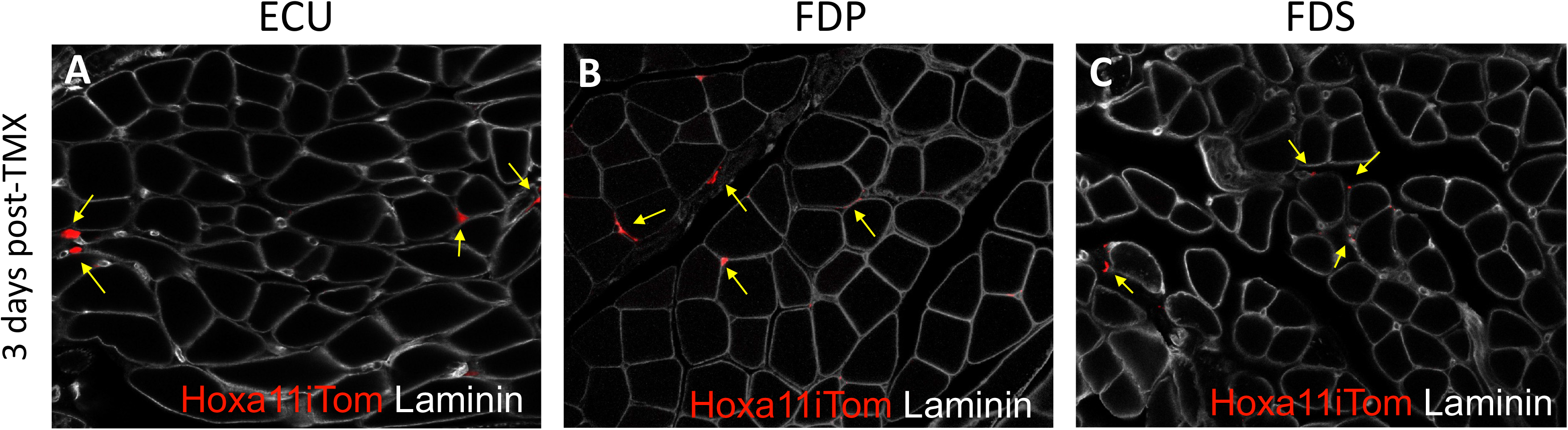
Hoxa11 expressing cells are only in the interstitium of zeugopod-attached muscles after initial recombination. Hoxa11 lineage labeling (Hoxa11iTom, red) is observed 3 days following tamoxifen administration. (**A-C**) Hoxa11 lineage-labeled cells are only seen in the interstitium (yellow arrows) of the Extensor Carpi Ulnaris (ECU), Flexor Digitorum Profundus (FDP) and Flexor Digitorum Sublimis (FDS) as shown by laminin (white).

**Figure S3.**
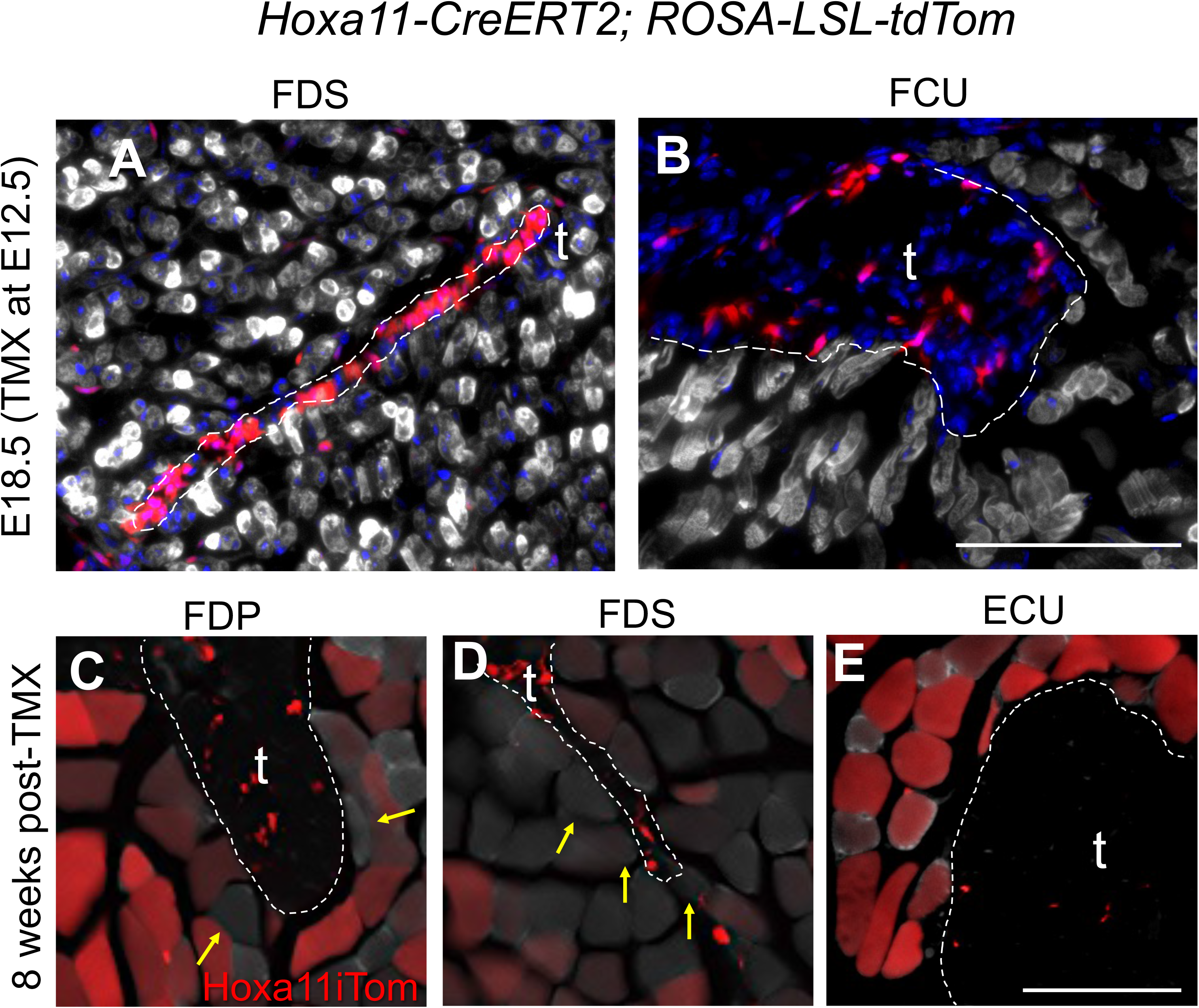
Hoxa11 lineage shows no preferential contribution to myofibers near the myotendinous junctions during embryogenesis or homeostasis. Sections of embryonic forelimb muscles were analyzed for myofiber lineage contribution at E18.5. (**A, B**) Images of the Flexor Digitorum Sublimis (FDS) and Flexor Carpi Ulnaris (FCU) muscles at the myotendinous junction show no Hoxa11iTom (red) overlap with My32 (white), consistent with lineage labeling throughout the rest of the muscle body. Sections from adult Hoxa11iTom muscle of the Flexor Digitorum Profundus (FDP), Flexor Digitorum Sublimis (FDS), and Extensor Carpi Ulnaris were analyzed for *Hox11* lineage contribution at the myotendinous junction 8 weeks after the induction of lineage reporting. High magnification images show similar lineage labeling near the myotendinous junction FDP (**C**), FDS (**D**), or ECU (**E**) as in the mid-body of the muscle (Figure 4). Scale bars = 100μm. Tendons are marked with a t.

**Figure S4.**
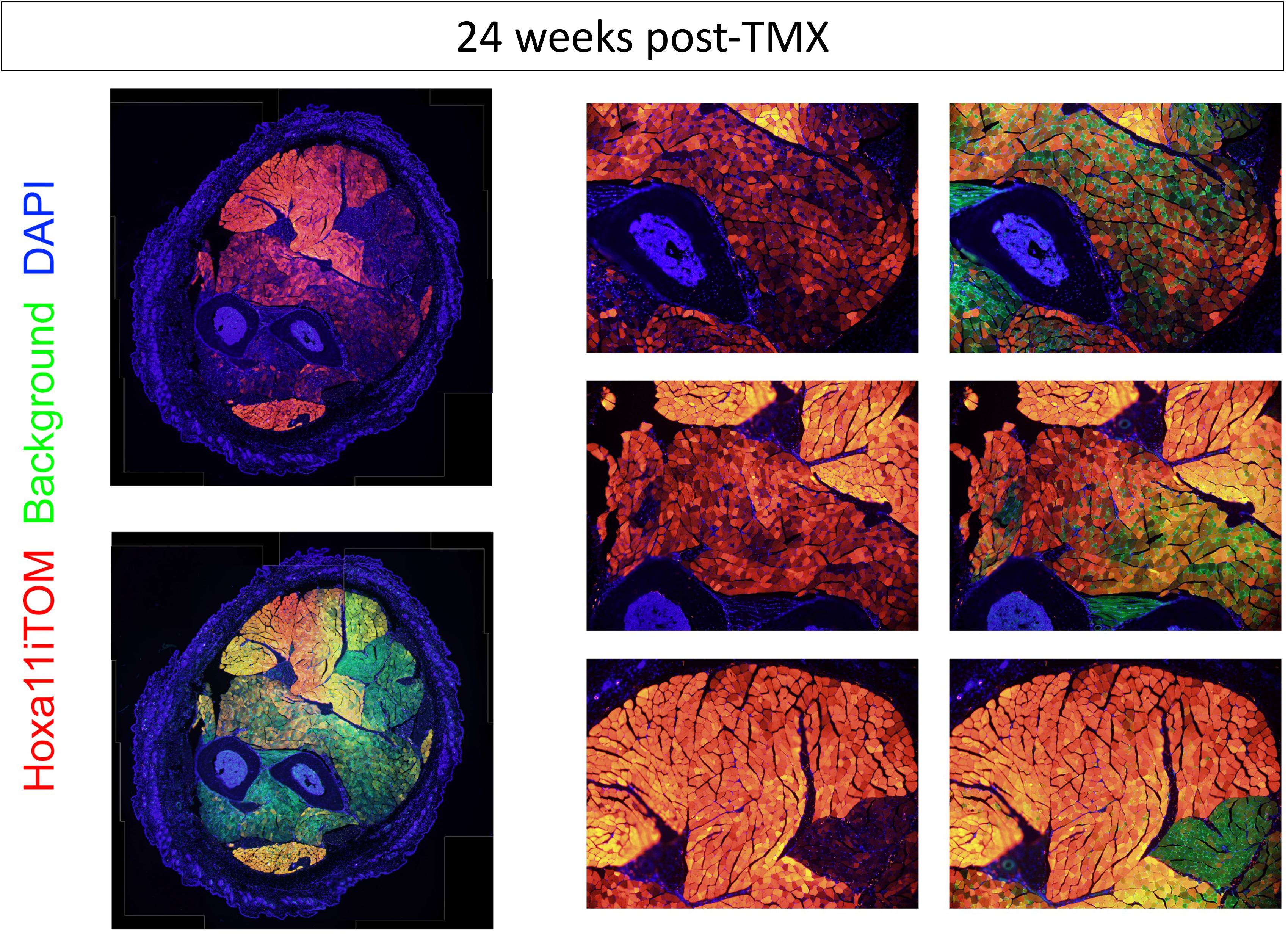
Hoxa11 lineage 24 weeks after reporter induction. Hoxa11^CreERT2/+^; ROSA^LSL-tdTomato/+^ mice were given 5mg tamoxifen at 8 weeks of age and collected 24 weeks (6 months) later. A full cross-section with and without background (green) show many Hoxa11iTom+ (red) muscle fibers. Close ups with and without background depict high levels of tdTomato within muscle fibers.

**Figure S5.**
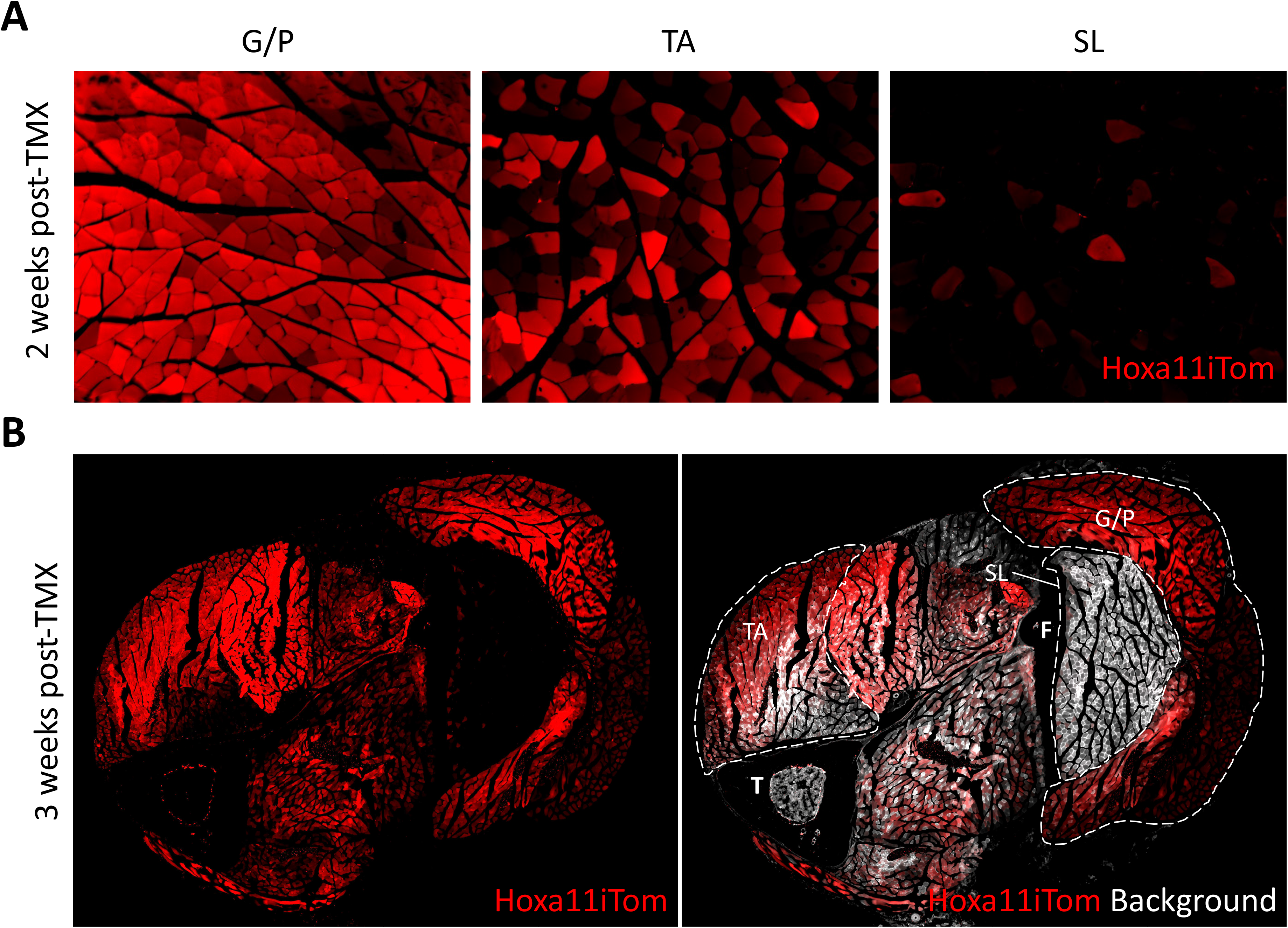
Hoxa11 lineage differentially contributes to hindlimb muscles. (**A**) Hindlimb muscles taken from Hoxa11iTom animals 2 weeks after tamoxifen treatment show different levels of lineage contribution in the Gastrocnemius/Plantaris (G/P), Tibialis Anterior (TA), and Soleus (SL). (**B**) A whole hindlimb cross-section from an animal 3 weeks after the start of lineage labeling shows variable tdTom expression in muscles of the hindlimb. T and F mark the tibia and fibula, respectively. Muscles shown in **A** are outlined and labeled in **B**.

**Figure S6.**
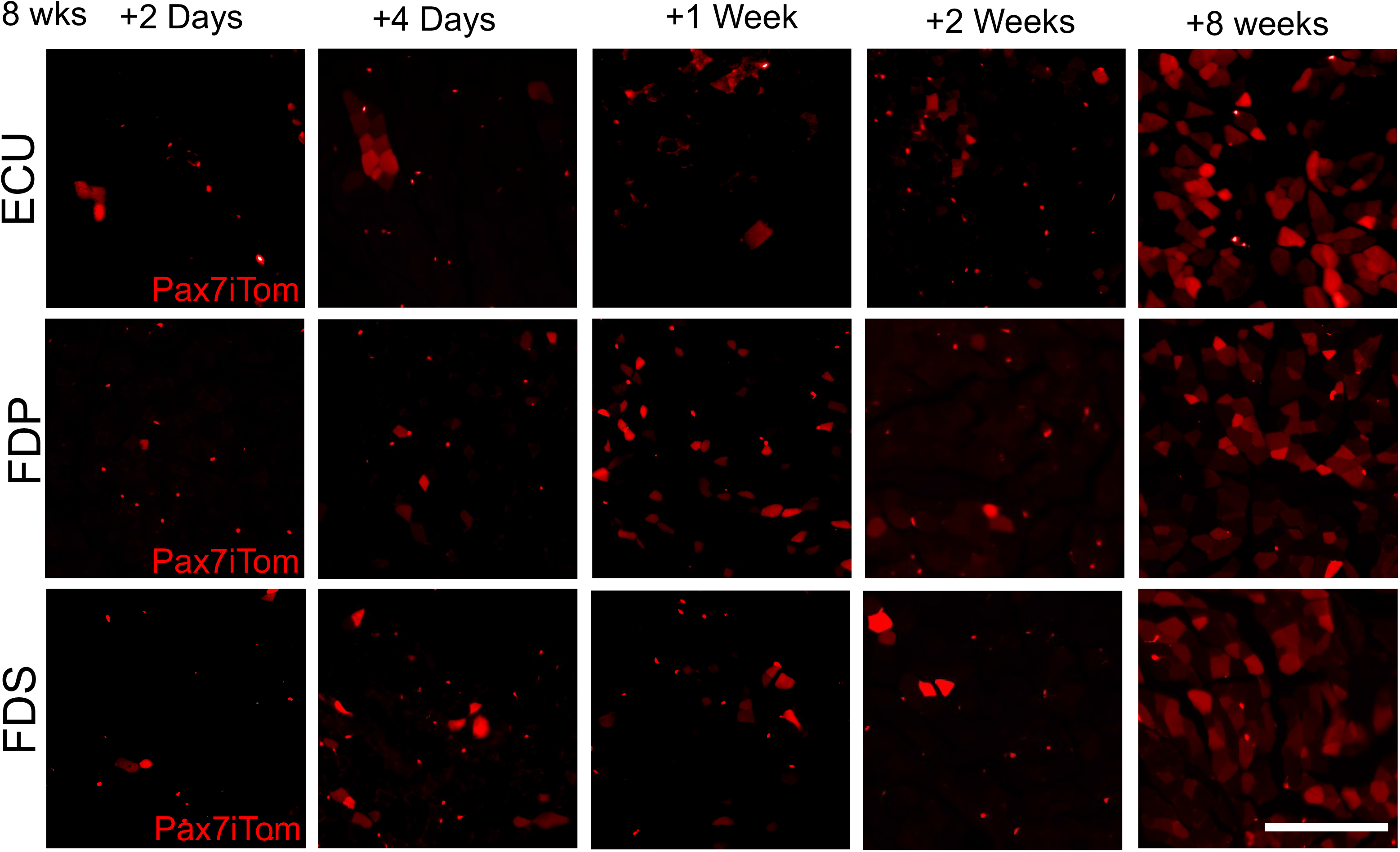
Pax7iTom lineage in the forelimb muscles. *Pax7CreERT2; ROSALSL-TdTomato* (Pax7iTom) mice were given a single intraperitoneal injection of 5 mg tamoxifen at 8 weeks of age and collected at 2 days, 4 days, 1 week, 2 weeks, and 8 weeks after tamoxifen induction. Images of the Extensor Carpi Ulnaris (ECU), Flexor Digitorum Profundus (FDP) and Flexor Digitorum Sublimis (FDS) muscles show tdtom+ myofibers as well as tdTom+ satellite cells.

**Figure S7.**
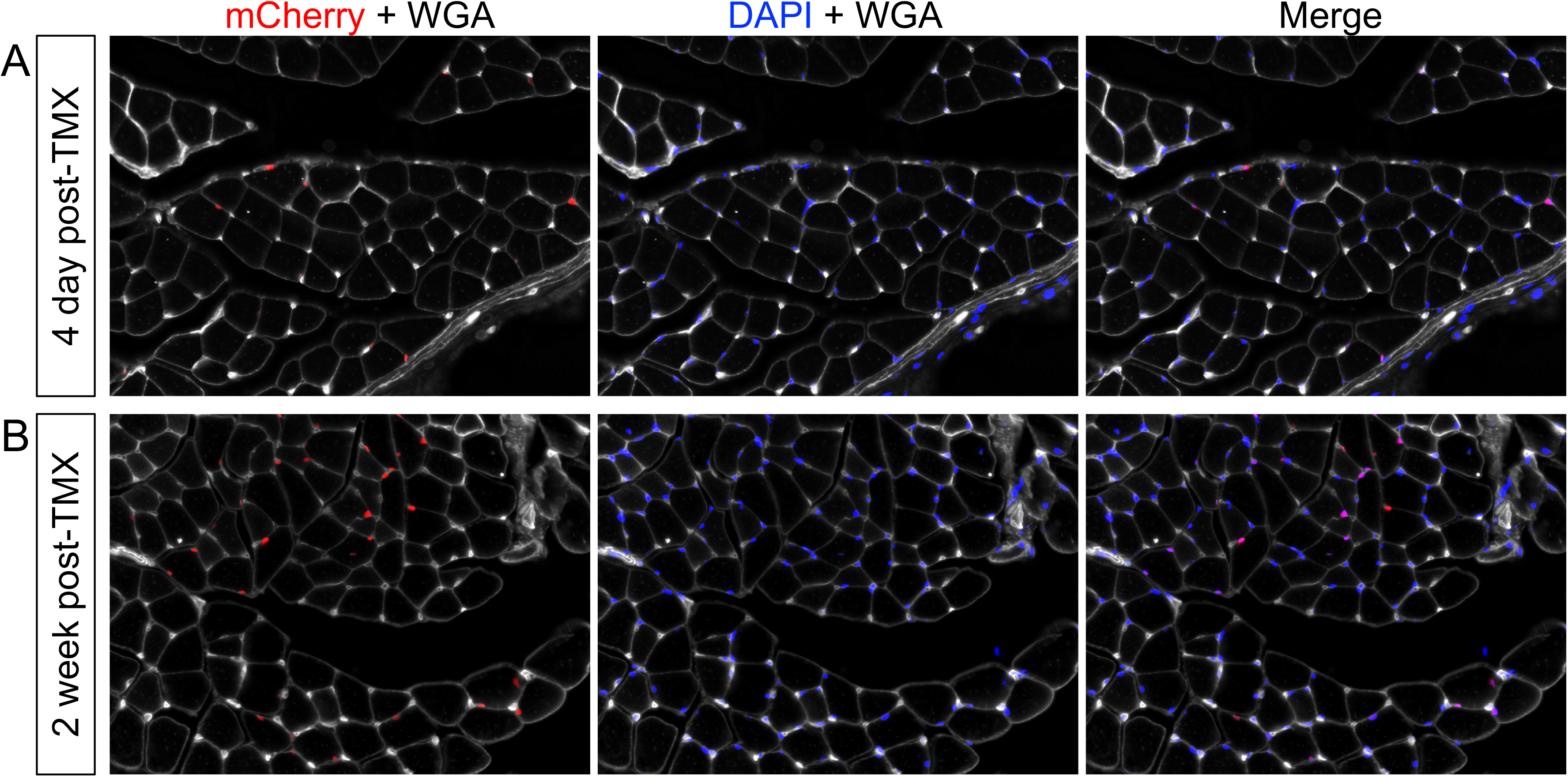
Hoxa11iH2BmCherry labeled nuclei overlap with nuclear stain. H2BmCherry expression in nuclei seen in figure 5 was confirmed with DAPI nuclear stain in images 4 days (**A**) and 2 weeks (**B**) post tamoxifen.

**Figure S8.**
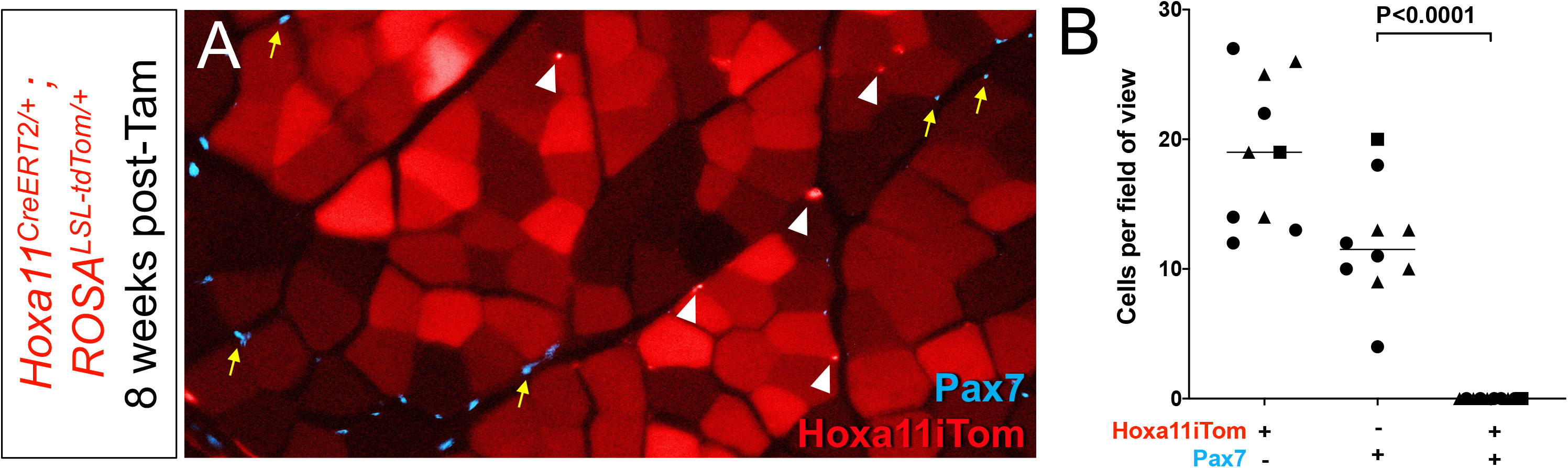
Hoxa11 lineage does label satellite cells 8 weeks post-induction of lineage reporter. (**A**) Forelimb muscle sections from animals 8 weeks (dosing at 8 weeks of age, collection at 16 weeks of age) after the start of lineage labeling were stained for Pax7 (cyan, yellow arrows) and analyzed for overlap of Hoxa11itom (red, arrowheads) and Pax7. (**B**) Quantification shows no Hoxa11iTom+/Pax7+ cells were observed through forelimb muscles.

**Table S1.** Antibodies.

| Name | Source | Catalog Number | Dilution | Antigen retrieval |
| --- | --- | --- | --- | --- |
| anti-GFP | Aves Labs | GFP-1010 | 1:200 | No |
| anti-Twist2 | Abcam | ab66031 | 1:200 | No |
| anti-PDGFRa | Abcam | ab61219 | 1:50 | No |
| anti-Tcf4 | Cell Signaling | 2569S | 1:100 | No |
| WGA-647 | Invitrogen | W32466 | 1:250 | No |
| WGA-488 | Invitrogen | W11261 | 1:250 | No |
| anti-Pax7 | DSHB | Pax7-S | 1:10 | Yes |
| anti-Laminin | Sigma | L9393 | 1:500 | No |
| anti-My32 | Sigma | M4276 | 1:400 | Yes |
| anti-PECAM1 | DSHB | 2H8-S | 1:10 | No |
| anti-BF-F3 | DSHB | BF-F3 | 1:10 | Yes |
| anti-SC-71 | DSHB | SC-71 | 1:10 | Yes |
| anti-Myosin (Skeletal, Slow) | Sigma | M8421 | 1:250 | Yes |
| anti-Scal(Ly6A/E)-PE | BD Pharmingen | 553108 | 1:100 | No |
| anti-Ter119-AF700 | BD-Pharmingen | 560508 | 1:100 | No |
| anti-CD31-PE-CF594 | BD-Horizon | 563616 | 1:100 | No |
| anti-ITGA7-AF647 | AbLab | 67-0010-05 | 1:100 | No |
| anti-CD45-AF700 | Invitrogen, eBioscience | 56-0451-80 | 1:100 | No |
| anti-CD106(VCAM1)-PE/Cy7 | Biolegend | 105719 | 1:40 | No |
| Alexa-488 Donkey anti-Chicken | Jackson ImmunoResearch Labs | 016-580-084 | 1:500 | No |
| Alexa-555 Donkey anti-Rabbit | Invitrogen | A31572 | 1:500 | No |
| Streptavidin-AF647 | Invitrogen | S21374 | 1:500 | No |

